# Peptide partitions and protein identification: a computational analysis

**DOI:** 10.1101/069526

**Authors:** G. Sampath

## Abstract

Peptide sequences from a proteome can be partitioned into N mutually exclusive sets and used to identify their parent proteins in a sequence database. This is illustrated with the human proteome (http://www.uniprot.org; id UP000005640), which is partitioned into eight subsets KZ*R, KZ*D, KZ*E, KZ*, Z*R, Z*D, Z*E, and Z*, where Z ∈ {A, N, C, Q, G, H, I, L, M, F, P, S, T, W, Y, V} and Z* ≡ 0 or more occurrences of Z. If the full peptide sequence is known then over 98% of the proteins in the proteome can be identified from such sequences. The rate exceeds 78% if the positions of four internal residue types are known. When the standard set of 20 amino acids is replaced with an alphabet of size four based on residue volume the identification rate exceeds 96%. In an information-theoretic sense this last result suggests that protein sequences effectively carry nearly the same amount of information as the exon sequences in the genome that code for them using an alphabet of size four. An appendix discusses possible *in vitro* methods to create peptide partitions and potential ways to sequence partitioned peptides.

## 1. Introduction

Partitioning of peptides into mutually exclusive subsets of sequences with known properties can make protein sequencing and/or identification based on database search easier. If a full or partial sequence from a subset is known identification of the parent protein is more efficient because the search space is considerably smaller. Here it is shown by computation that a partition with N = 8 sets can lead to protein identification rates in excess of 90%. An appendix discusses practical methods for *in vitro* cleaving of proteins, physical separation of cleaved peptides into subsets, and full or partial sequencing of partitioned peptides.

## 2. Peptide partitions and their properties

Consider peptide sequences in a proteome that have the form X_1_Z*X_2_, where X_1_ and X_2_ are drawn from a small number of residue types, Z is one of the remaining residue types, and Z* ≡ 0 or more occurrences of Z. Table 1 shows three such partitions for different X_1_ and X_2_ at some pH value.

**Table 1.**
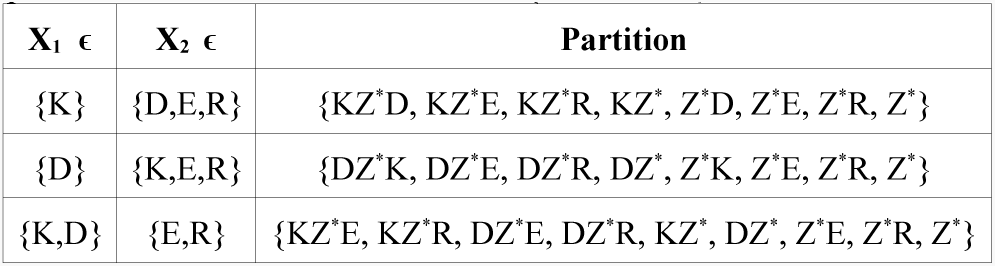
Three partitions of the form X_1_Z*X_2_; Z ∈ {A, N, C, Q, G, H, I, L, M, F, P, S, T, W, Y, V}

| $X_1 \in$ | $X_2 \in$ | Partition |
| --- | --- | --- |
| {K} | {D,E,R} | {$KZ^*D$, $KZ^*E$, $KZ^*R$, $KZ^*$, $Z^*D$, $Z^*E$, $Z^*R$, $Z^*$} |
| {D} | {K,E,R} | {$DZ^*K$, $DZ^*E$, $DZ^*R$, $DZ^*$, $Z^*K$, $Z^*E$, $Z^*R$, $Z^*$} |
| {K,D} | {E,R} | {$KZ^*E$, $KZ^*R$, $DZ^*E$, $DZ^*R$, $KZ^*$, $DZ^*$, $Z^*E$, $Z^*R$, $Z^*$} |

The following are some properties of such partitions; they are derived in part from computations performed on the complete sequence database of the human proteome (20207 sequences).

1. Partitioning makes protein identification based on comparison of peptide sequences with sequences in a proteome database simpler because it reduces the search space considerably. This can be seen in Table 2, which shows the number of peptides in each subset of an 8-set partition and the computed number of protein-identifying peptides in it; the average reduction of the search space is around 8. Also, on computing the set union operation (U) over the identifying peptides in different subsets the percentage of identifiable proteins in a proteome increases appreciably. For example with KZ*R U KZ*D U KZ*E the number exceeds 90%. With all sets the rate exceeds 98%. Note that with Z* an element can be a protein identifier and at the same time be a subsequence of a peptide in any of the other seven subsets. This is because it only needs to be a unique identifier for a protein containing an element of Z*; being in the distinct subset Z*it carries context information that is hidden. Thus in the protein sequence that it identifies it is not preceded by K in any sequence in the proteome, it is either followed by K or is the last peptide at the end of the protein sequence, and it is not followed by R, D, or E.
2. With partitioned peptides, sequencing of peptides in one subset is not affected by peptides in another because they occur in two distinct and physically separate sequencing procedures. For example, sequencing of peptides in KZ*R is independent of sequencing in KZ*D. Thus peptides in the two subsets can be sequenced in parallel and the results logically combined.
3. Sequences in {KZ*R}, {KZ*D}, and {KZ*E} do not have internal residues from {K,R,D,E}. When sequencing a peptide in any of these subsets its internal sequence need be checked against 16 rather than 20 residue types. Similar statements can be made about the other subsets.
4. In most cases, when sequencing a peptide a small number k of internal residue types may be sufficient to identify it and its container protein. With k = 2, 3, and 4 there are 120, 560, and 1820 ways to choose residue types. Figure 1 shows the number of proteins in the human proteome (20207 proteins) that are identifiable from partial sequences in KZ*R, KZ*D, and KZ*E for k = 2 and 4. (Residue types are selected in the order of their frequency of occurrence in the proteome; see Figure A-5 in the Appendix.) A set union operation further increases the total number of identifiable proteins. The maximum number identifiable can be obtained by exhaustive search through all 120, 560, or 1820 tagging choices with k = 2, 3, or 4; it can be expected to exceed 78% of the total number of proteins.

**Figure 1.**
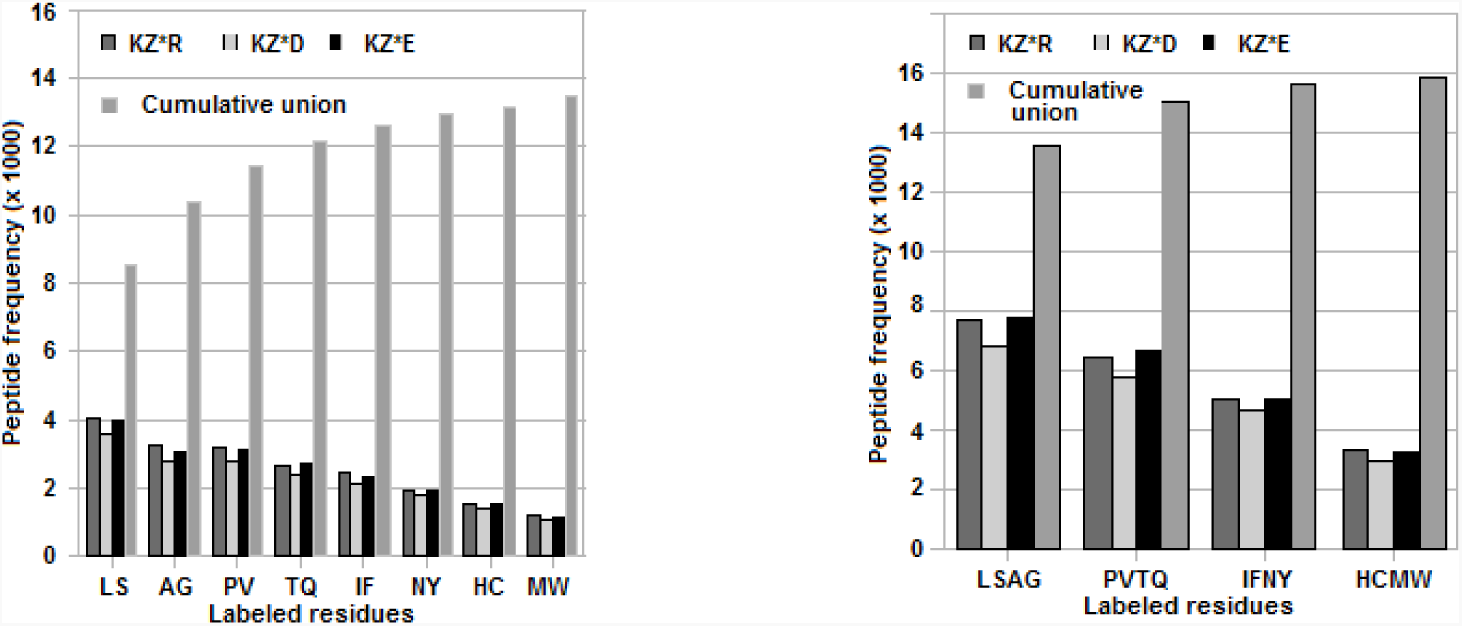
Number of peptides in KZ*R, KZ*D, and KZ*E in the human proteome identifiable from two (left panel) or four (right panel) internal residues. The fourth bar in each group is the frequency of cumulative unions from LS to MW or from LSAG to HCMW. Thus the rightmost bar in the two cases is the percentage total of identifiable peptides in the proteome (66%, left panel; 78%, right panel).

**Figure A-5.**
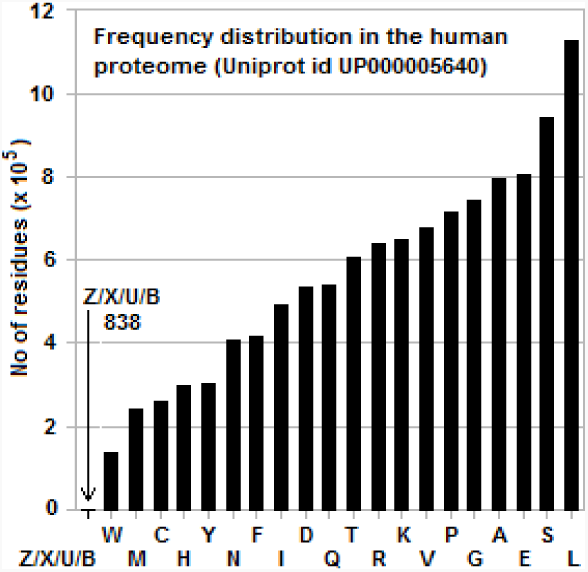
Frequency distribution of residues in the human proteome (Uniprot id UP000005640) (Z/X/U/B represents 838 ambiguously labeled residues in the sequence database.)

**Table 2.** Partitioned subset sizes for the human proteome (Uniprot database id UP000005640; 20207 sequences) and protein identification efficiency (column 4 ÷ 20207)*100 of full peptide sequences. (Last row: U = union.)

| Partition subset | Total no. of peptides | No. of protein-identifying peptides | No of proteins identified | Percentage identified |
| --- | --- | --- | --- | --- |
| $KZ^*R$ | 139423 | 38110 | 14247 | 70.51 |
| $KZ^*D$ | 125351 | 34830 | 12983 | 64.25 |
| $KZ^*E$ | 194024 | 45440 | 14304 | 70.79 |
| $KZ^*$ | 189713 | 44067 | 13736 | 67.98 |
| $Z^*R$ | 499784 | 128902 | 18305 | 90.59 |
| $Z^*D$ | 411189 | 110708 | 17691 | 87.55 |
| $Z^*E$ | 609872 | 140051 | 18254 | 90.34 |
| $Z^*$ | 345450 | 111045 | 18467 | 91.39 |
| $KZ^*R \cup KZ^*D \cup KZ^*E$ | 458798 | 54309 | 18212 | 90.13 |
| Union of all 8 subsets | 2514806 | 287809 | 19885 | 98.4 |

## 3. A partition based on a reduced amino acid alphabet

Amino acids vary in volume [1] but in most cases the differences are small. In [2] they are coarsely partitioned into four subsets labeled *Minuscule*, *Small*, *Intermediate*, and *Large*. These subsets are encoded here with the reduced alphabet {B, U, X, Z}. (This alphabet, which is not part of the 1-letter standard amino acid code, was chosen to make the database search algorithms simpler.) The mapping is as follows: {G,A,S,C} → B, {T,D,P,N,V} → U, {E,Q,L,I,H,M,K} → X, {R,F,Y,W} → Z. The sequences in the proteome database are recoded with this reduced alphabet and protein identifying peptides are obtained by searching for them in the sequence database.

By redefining the subsets three more volume-based partitions with 5 or 6 subsets are possible: 1) {G} → {G} {A,S,C} → B, {T,D,P,N,V} → U, {E,Q,L,I,H,M,K} → X, {R,F,Y,W} → Z; 2) {G,A,S,C} → B, {T,D,P,N,V} → U, {E,Q,L,I,H,M,K} → X, {R,F,Y} → Z, {W} → {W}; and 3) {G} → {G}, {A,S,C} → B, {T,D,P,N,V} → U, {E,Q,L,I,H,M,K} → X, {R,F,Y} → Z, {W} → {W}. Table 3 presents two sets of results for the 8 subsets of the peptide partition {Z*D, Z*E, KZ*D, KZ*E, Z*, KZ*, Z*R, KZ*R} using the reduced codes {B, U, X, Z} and {G, B, U, X, Z, W}. (Compare these results with Figure 1, which shows the identification efficiency of partitions KZ*R, KZ*D, and KZ*E in which the positions of 4 or 6 residue types in a peptide are known.)

**Table 3.** Calculated number of peptides in peptide sequences coded with reduced amino acid alphabets identifying container protein and protein identification efficiency in human proteome (Uniprot database id UP000005640; 20207 sequences)

| Partition subset | No of peptides | Reduced amino acid alphabet |  |  |  |  |  |
| --- | --- | --- | --- | --- | --- | --- | --- |
|  |  | {B, U, X, Z} |  |  | {G, B, U, X, Z, W} |  |  |
|  |  | No. of identifying peptides<sup>(a)</sup> | No. of uniquely id'd proteins<sup>(b)</sup> | Identification efficiency<sup>(c)</sup> | No. of identifying peptides<sup>(a)</sup> | No. of uniquely id'd proteins<sup>(b)</sup> | Identification efficiency<sup>(c)</sup> |
| KZ*R | 139423 | 18097 | 10060 | 49.78% | 21976 | 11215 | 55.5% |
| KZ*D | 125351 | 16267 | 9019 | 44.63% | 19785 | 10048 | 49.72% |
| KZ*E | 194024 | 19945 | 10185 | 50.4% | 24291 | 11270 | 55.77% |
| KZ* | 189713 | 19737 | 9832 | 48.65% | 24077 | 10877 | 53.82% |
| Z*R | 499784 | 57937 | 15770 | 78.04% | 72752 | 16642 | 82.35% |
| Z*D | 411189 | 48944 | 14700 | 72.74% | 61304 | 15673 | 77.56% |
| Z*E | 609872 | 58914 | 15690 | 77.64% | 74552 | 16612 | 82.2% |
| Z* | 345450 | 47968 | 15370 | 76.06% | 59570 | 16423 | 81.27% |
| KZ*R U KZ*D U KZ*E | 458798 | 54309 | 15744 | 77.91% | 66052 | 17139 | 84.81% |
| Union of all 8 sets | 2514806 | 287809 | 19581 | 96.9% |  |  |  |

<sup>(a)</sup> Number of peptides in partition subset that can uniquely identify their container protein
<sup>(b)</sup> Total number of proteins in proteome uniquely identified by the identifying peptides in column marked <sup>(a)</sup>
<sup>(c)</sup> Protein identification efficiency = number in column marked <sup>(b)</sup> \* 100/20207

As an example, consider the 8 partitioned subsets discussed earlier, namely Z*D, Z*E, KZ*D, KZ*E, Z*, KZ*, Z*R, and KZ*R. There are a total of 2,514,806 peptides in the full partition. Consider one of the subsets, KZ*R. It has 139423 peptides. When recoded with the reduced alphabet 18097 of them can identify their parent protein. The number of unique proteins identified is 10060, which is 49.78% of the 20207 proteins in the human proteome. (With the full alphabet, the corresponding numbers are 38110, 14247, and 70.51%.) When the results from different subsets are subjected to the union operation the identification rate increases significantly. For example, the identification efficiency of KZ*R U KZ*D U KZ*E with a 4-character alphabet is 77.91%, while with the union of all eight sets it is 96.9%. (Further post-processing using HMM or similar learning-based algorithms [3-6] can result in further improvements in identification, but such a development is outside the scope of the present report.)

These results show that almost all (> 96%) proteins in the proteome can be identified when they are coded with a reduced four-character alphabet. This suggests that at the identification level protein sequences contain, in an information-theoretic sense, about the same amount of information as the exons in the genome that code for them. Whether this is biologically significant or is a computational artifact remains to be seen.

## Appendix

### Contents

A-1 Peptide partitioning *in vitro*

A-2 Peptide sequencing in practice

A-3 Frequency distribution of residues in the human proteome

#### A-1 Peptide partitioning in vitro

Partitioning of peptides can be done *in vitro* in two steps: cleaving and separation. High-specificity peptidases or chemical agents are available to cleave before or after a specific residue [7,8], where ‘before’ or ‘after’ means cleavage on the N-terminal or C-terminal side of the residue. (Peptide sequences are usually listed from N-terminal to C-terminal.) Some of these peptidases/agents are listed in Table A-1 below. Peptides can be separated using a variety of methods based on the electric charge they carry. (This charge can be calculated using the Henderson-Hasselbach equation [9].) Examples of separation methods [10] include isoelectric focusing (IEF), paper chromatography (PC), and ion-exchange chromatography (IEC).

**Table A-1:**
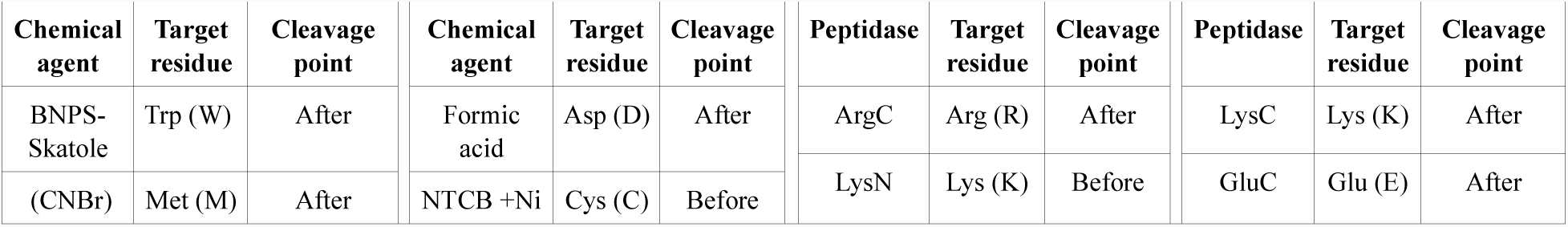
Selected peptidases/chemicals, their targets, and where cleavage occurs

In Step 1 proteins in the assay sample are broken into peptides using multiple peptidases or chemical agents in sequence [11]. This enables breaking a protein into peptides with desired start and end residues (see Table 1 in the main text). For example LysN creates peptides starting with K, while formic acid creates peptides ending in D. When these two are applied in sequence (in any order), the result is peptides that start with K and end with D. With four peptidases/chemical agents, there are 24 different orders of application. The optimum order may be determined experimentally.

In Step 2 the peptides created in Step 1 are inserted into an electrolyte with a known pH value and separated into subsets based on the charge they carry at that pH value. IEF uses an electrolyte with a pH gradient and an electrical field that moves a peptide to a point along the gradient where its effective charge is zero. This point is the peptide’s ‘isoelectric focus’ or pI value. IEF has a pI resolution of 0.001.

Figure A-1 is a flow chart that shows cleavage of proteins into peptides of the form Z*D, Z*E, KZ*D, KZ*E, Z*, KZ*, Z*R, and KZ*R, followed by IEF. This leads to an 8-subset partition in which the maximum peptide frequency in each subset is separated from that in another by a pI value of at least 0.1. For the human proteome the distribution of peptide frequency vs pI for each of the 8 subsets of the partition is shown in Figure A-2.

**Figure A-1.**
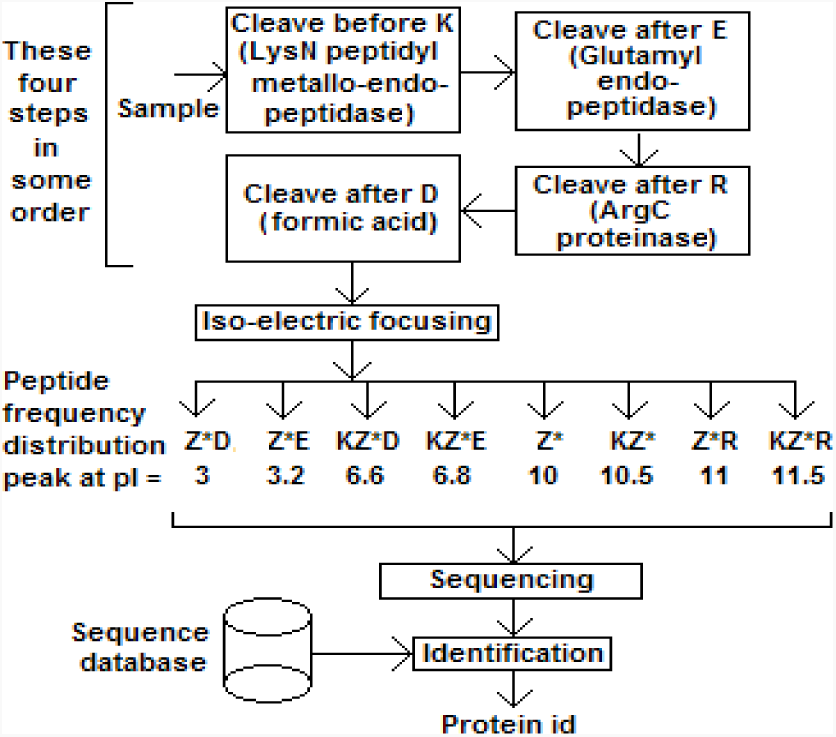
Flow chart for *in vitro* separation of peptides into 8 subsets (followed by sequencing and protein identification).

**Figure A-2.**
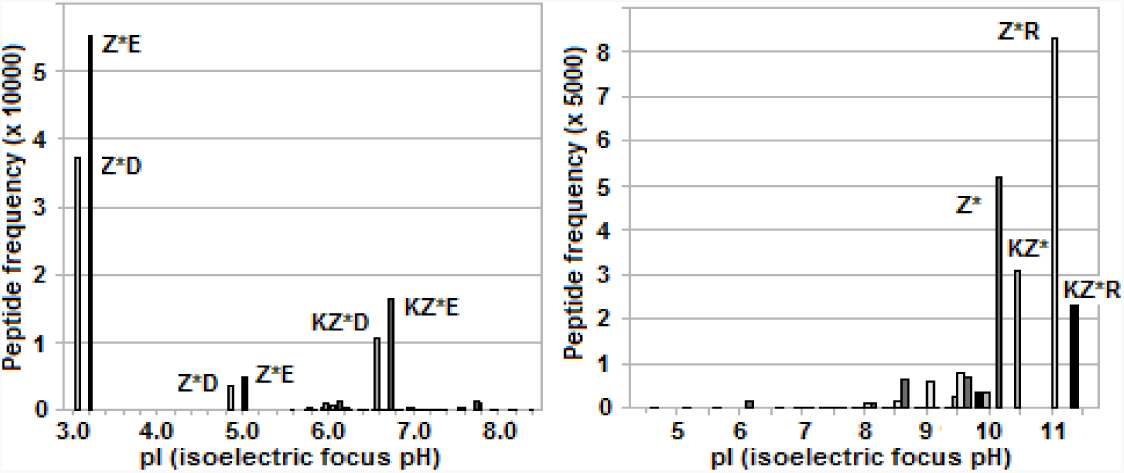
Frequency distribution of peptides in 8-subset partition of human proteome vs pI (isoelectric focus pH).

Peptides can be further separated by mass or size using 2-D gel electrophoresis (GE) [12]. Figure A-3 shows a somewhat oversimplified example of a computed distribution of peptides in {KZ*R} by mass that resembles analyte separation in the vertical dimension in 2-D GE.

**Figure A-3.**
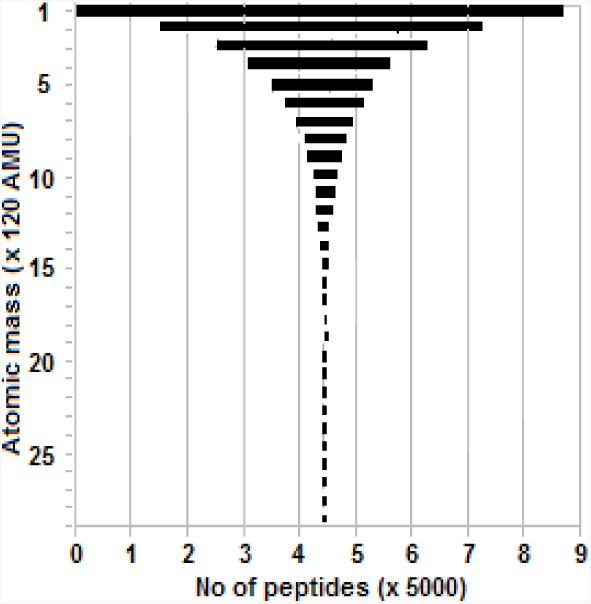
Distribution of peptides in {KZ*R} by mass. (Horizontal bars manually centered.)

In practice each step before sequencing in Figure A-1 requires an extraction step involving elution, filtration, etc., in which the target (peptide fragments) is extracted and input to the next step. With the 8-subset partition example considered above five extraction steps are required, one after each cleavage step and one after IEF. This means that even if each extraction is 90% efficient, less than 60% of the sample will enter the sequencing stage.

#### A-2 Peptide sequencing in practice

Two practical methods to sequence peptides in partitioned subsets are considered here. They are mass spectrometry (MS) [13], which sequences peptides in the bulk, and nanopore sequencing, which is a single molecule method [14,15].

In MS, peptides are extracted from the gel after GE and split into ionized fragments before entering the spectrometer, which then outputs a mass spectrum for a peptide. A variety of algorithms are available to predict a partial or full sequence from the spectrum by comparison with MS spectra databases and identify the container protein [16,17]. With partitioned peptides, the task is simplified because a peptide belongs to a restricted subset.

In conventional nanopore sequencing of polymers (DNA, RNA, protein) the analyte translocates through the pore by a combination of diffusion and electrophoresis in the presence of an electric field. Discrimination among monomers (bases in DNA and RNA; residues in peptides) is based on the ability to detect measurable differences in the current blockade due to different monomers. Protein sequencing is more difficult than DNA sequencing because of the need to discriminate among 20 residue types against 4 base types. This is further complicated by homopolymers because successive blockade levels have the same value. One way to resolve this is to use a pore in an atom-thin membrane of graphene or MoS_2_. The analyte translocation distance through the pore is then less than the distance between two successive monomers so blockade transitions between two monomers can be better distinguished.

Sequencing that is based on pore current blockades has to contend with ever present noise, so a high signal to noise ratio (SNR) is required. One way to increase SNR is to increase the transmembrane potential and thereby increase pore current levels; however it cannot exceed a few 100s of millivolts. Additionally the analyte takes less than 100 ns to pass through the pore, which puts detection beyond the capacity of available detectors. Even with available detector bandwidths, higher bandwidth means higher noise levels. Several methods of slowing down the analyte have been proposed [18], most of them fall short. Two potential exceptions to this are: 1) slowdown based on the use of a hydraulic pressure gradient that is opposed to the transmembrane potential [19]; and 2) use of a high viscosity electrolyte (= RTIL or room temperature ionic liquid) [20] in the *cis* chamber, which has been found to decrease translocation speeds by a factor of ∼200 in DNA sequencing with a pore in an atom-thin MoS_2_ membrane.

In [2] peptide sequencing is based on using a reduced amino acid alphabet with four characters and mapping current blockades caused by residues to it. Post-processing of signal data with four measurable levels of discrimination uses a hidden Markov model (HMM) to construct the reverse map from reduced alphabet to full alphabet and the peptide sequence therefrom. If the notion of a reduced alphabet is combined with that of peptide partitions, nanopore sequencing can in principle identify more than 96% of the proteins in the human proteome (see Table 3 in the main text).

The dependence on pore currents to detect monomers can be avoided if selected residue types in a peptide are labeled with fluorescent tags and detected optically using Total Internal Reflection Fluoroscopy (TIRF) [21,22]. This results in a partial sequence for a peptide, which can be used to identify the parent protein through database search. (Pore current blockades, while still present, are not used for monomer detection.) Figure A-4 shows a schematic. Peptide translocation through the pore is controlled by a combination of transmembrane potential and hydraulic gradient, as in [19], and/or by using an RTIL, as in [20]. As the peptide threads through the pore into *trans* an emerging residue that is tagged fluoresces in the presence of a laser input and is identified by a photosensor. With additional signal processing positional information for the tagged residues in the peptide sequence is obtained.

**Figure A-4.**
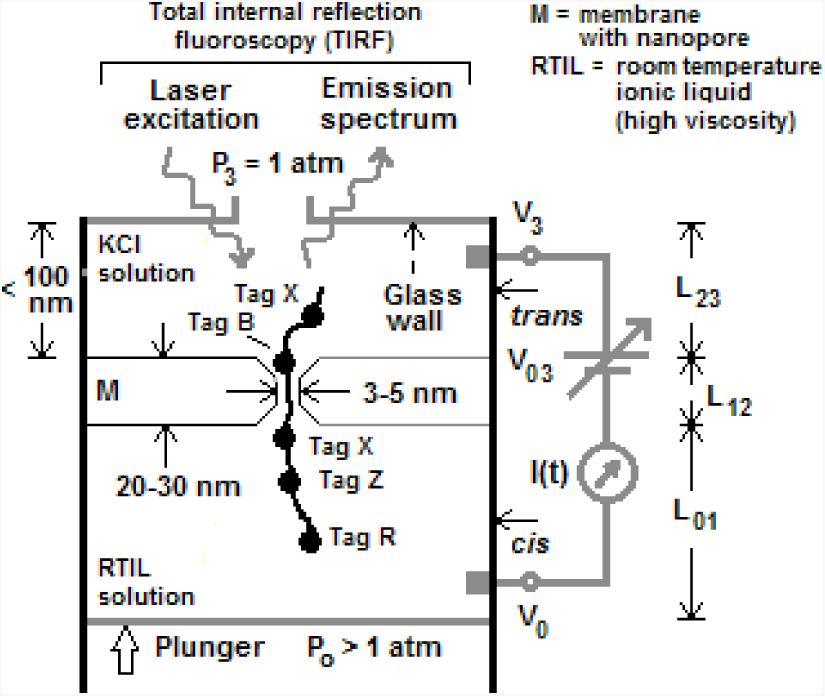
Schematic of cell with hydraulic gradient, transmembrane potential, RTIL in *cis*, and TIRF. (Not to scale) (Peptide cartoon shows element of Z*R with 4 tag types: one for end residue R and three for internal residue types Z, X, and B.)

TIRF has a resolution in the submillisecond range, which appears achievable with the above setup. It can distinguish up to 13 intensity levels so peptides can contain up to 13 tagged residues of the same type. It has the further advantage of being able to sense homopolymers correctly because every residue in the homopolymer is detected by a quantum jump in the photo-intensity, unlike with electrical sensing in which pore current measurements yield blockade current levels that are the same for successive identical residues.

Consider for example sequencing peptides in KZ*R, KZ*D, and KZ*E. Residues K, R, E, and D can be coded using two tags, denoted TERM-1 and TERM-2 (K and R; K and E; K and D). A small number k of internal residue types from the set Z can also be tagged. With k = 2 let the latter be INTL-3 and INTL-4. A tagged residue causes a step increase in the intensity of the measured signal TERM-m(t) or INTL-n(t) from the nanopore for tag type m or n. The residue order can then be constructed from the individual signals TERM-1(t), INTL-3(t), INTL-4(t), and TERM-2(t). Position information can be estimated from time differences between successive tag detection times obtained from the signals TERM-m(t) and INTL-n(t) using deconvolution algorithms similar to those used in DNA sequencing [3-6]. From the resulting partial sequence the full peptide sequence can be identified by database search. If it is also a unique identifier for its container protein the latter can be identified as well. Figure 1 in the main text shows calculated frequency distributions of identifying peptides in KZ*R, KZ*D, and KZ*E with two or four internal tag types.

With KZ*R, KZ*D, and KZ*E, the boundary between two peptides that pass through the pore in succession is known from the labels of the first and last residues. Whether the peptides enter C-terminal or N-terminal first, their end labels always occur in pairs (for example, with labeled internal residues X and Z: K.X.Z..R, K.Z…Z…..R, RXX..Z…K, etc.). With KZ*, Z*R, Z*D, and Z*E, only one starting/terminal residue is fixed. With a total of four tag types, the signals for KZ* would be TERM-1(t), INTL-2(t), INTL-3(t), and INTL-4(t). With Z* there is neither a starting nor an ending label, so the first signal is replaced by INTL-1(t). Two successive peptides S_1_ and S_2_ are detected as follows:

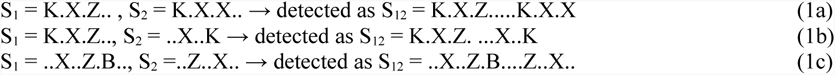

In each case the boundary between the two peptides can be discerned by searching for partial sequences S_1_’ and S_2_’ such that S_1_“..S_2_” = the observed S_12_, where S_1_“ (or S_2_”) = S_1_’ (or S_2_’) or its reverse. As an example Table A-2 shows the computed number of identifiable proteins from peptides with a total of four tag types whose partial sequences are truncated to the first tagged residue at one or both ends before search. For example, K.X.Z… is truncated to K.X.Z and ..X..Z.B.. to X..Z.B, while K.Z…Z…..R needs no truncation.

**Table A-2.** No of identifying peptides in three partitioned subsets of the human proteome with start/end and internal residues tagged, with and without truncation at one or both ends. (No truncation with KZ*R.)

| Property ↓ | Subset→ | KZ*R | Z*R | Z* |
| --- | --- | --- | --- | --- |
| No. of peptides identifying a protein |  | 14247 | 18305 | 18467 |
| Tagged residues |  | K L S R | L S A R | L S A G |
| Tagged peptides identifying protein |  | 4053 | 11560 | 12882 |
| With end(s) truncated before search |  | - | 10668 | 11398 |

Nanopore-based peptide sequencing can also be used to quantify proteins in an assay sample. Consider a mixture of proteins {(N_i_, P_i_, I_i_): i = 1,2, …} where Ni is the number of molecules of the i-th protein in the mixture, Pi the number of peptides per molecule of the protein (= the number of peptides created from a single molecule by the cleaving steps in Figure 1, main text), and I_i_ (0 ≤ I_i_ ≤ P_i_) the number of identifying peptides per molecule (this is known by precomputation; see Table 2, main text). Let N_total_ be the total number of peptides sensed and I_i-identified_ the number of identified peptides in protein i. If peptide entry into a cell is totally random, then after a sufficiently long run N_i_ can be estimated as 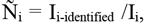, the corresponding fraction is 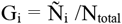. The number of peptides that do not yield identifying information is 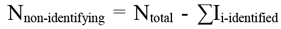, where the summation is over all the identified proteins. This number includes peptides that are found in more than one protein as well as impurities in the input sample.

#### A-3 Frequency distribution of residues in the human proteome

The 20 residue types occur with different frequencies in the human proteome. Figure A-5 shows their distribution. Strategies for peptide sequencing and protein identification may take this into account.

